# Microcystins are critical for the toxic *Microcystis* to survive long-term nitrogen starvation

**DOI:** 10.1101/2024.08.03.606466

**Authors:** Xiao-Ya Lian, Guo-Wei Qiu, Wen-Can Zheng, Jin-Long Shang, Hai-Feng Xu, Guo-Zheng Dai, Nan-Qin Gan, Zhong-Chun Zhang, Bao-Sheng Qiu

## Abstract

Toxic cyanobacterial blooms have expanded and intensified on a global scale, but the physiological role of microcystins during bloom development is not fully resolved. Here, we show that microcystin production can increase the survival and resuscitation rate of *Microcystis* after long-term nitrogen starvation. Our results showed that microcystin production could enable toxic *Microcystis* to accumulate more carbon reserves under nitrogen limitation, which is critical to support the survival of cells under stressful conditions. Further analysis showed that genes involved in microcystin synthesis were significantly upregulated at the initial phase of recovery, which could help toxic *Microcystis* to strengthen glycogen catabolism and fuel recovery. The close genetic traits between *Microcystis* strains suggest the strategies observed here might be highly conserved. Our findings imply how toxic *Microcystis* establish a competitive advantage over non-toxic species and provide new insight into the seasonal dynamic of the *Microcystis* population in natural environment.

**IMPORTANCE:** Microcystins are the most abundant cyanotoxins released during harmful algal blooms. While the factors controlling microcystin production have been widely studied, the function of these toxic secondary metabolites under changing environments remains poorly understood. Here we proved that microcystins are critical to toxic *Microcystis* to maintaining carbon metabolism under long-term nitrogen starvation and subsequent recovery. Compared to the non-toxic strains, microcystin-producing *Microcystis* exhibit a higher viability and resuscitation rate after prolonged nitrogen starvation, which is consistent with the dominance of these species at the early stage of cyanobacterial blooms. Our findings shed light on the genetic traits that drive population succession during bloom development, which is important for the modeling and prediction of harmful cyanobacterial blooms.

## INTRODUCTION

Harmful algal blooms are a major environmental problem worldwide (1, 2). It deteriorates water quality by depleting dissolved oxygen and producing toxic compounds, seriously impairing ecosystem function and threatening public health (3, 4). Recent studies have demonstrated that the frequency, intensity, and duration of cyanobacterial blooms are increasing on a global scale (1, 5–7). However, while the environmental drivers promoting harmful cyanobacterial blooms have been widely investigated, the mechanisms that bloom-forming cyanobacteria establish a competitive advantage over other species during bloom development were not fully understood. Many bloom-forming cyanobacteria are known to produce toxic secondary metabolites termed cyanotoxins (1, 8). Amongst them, microcystins (MCs) represent the most diverse and abundant families. MCs are cyclic non-ribosomal heptapeptides produced by *Microcystis*, *Planktothrix*, *Oscillatoria*, and *Dolichospermum* (9). It can cause severe liver and kidney damage when ingested by mammals and humans but also has harmful effects on neuro and reproduction (1). Although recent studies highlight the role of MCs in defending against grazing (10, 11), phylogenetic analysis reveals that their biosynthesis genes likely evolved before the origin of metazoan predators, indicating that the additional physiological function of MCs may exist (12, 13).

It has been suggested that the capacity of toxic cyanobacteria to synthesize MC is tightly associated with growth conditions (14–19). Among the environmental factors affecting MC production, nitrogen (N) availability is the most important one (20, 21). Firstly, MCs are N-rich products that contain a number of unusual amino acid residues. The N:C ratio of MC-LR is 10:49, which is much higher than the Redfield ratio of phytoplankton (16:106). Thus, it was not surprising that increased nitrogen concentration could promote MC production, whereas nitrogen depletion decreased MC synthesis (21, 22). Secondly, laboratory studies on *Microcystis aeruginosa* PCC 7806 (hereafter *Microcystis* 7806) revealed that the expression of MC biosynthetic genes (*mcy*) was regulated by the global nitrogen regulator NtcA, indicating a direct control of N availability in MC production (23, 24). Moreover, long-term field observations showed that the currently rising N:P ratio and increased N loading may favor non-nitrogen-fixing *Microcystis* spp. and MCs production (20, 25, 26). However, despite the high dependency of MC production on the nitrogen supply, the N bioavailability in surface waters frequently varies seasonally (20, 27, 28), and the physiological role of MCs under N fluctuation was largely unknown.

Here, we systematically investigated the physiological response of the toxic *Microcystis* and its MC biosynthesis deficient mutant to long-term N starvation and subsequent recovery. Our results show that MC production increases the growth and viability of this non-diazotrophic bloom-forming cyanobacteria in N-fluctuating water. The presence of MC-producing pathways could provide toxic cyanobacteria a competitive advantage over no-toxic strains, which enables them to quickly recover and develop dense blooms when combined N are plentiful.

## MATERIALS AND METHODS

### Culture conditions and general methods

*Microcystis aeruginosa* PCC 7806 (WT) and its *mcyB* mutant were provided by the Pasteur Culture Collection of Cyanobacteria (PCC, France). The successful insertion of the chloramphenicol-resistant fragment into the *mcyB* gene was confirmed by PCR (Fig. S1A). High performance liquid chromatography (HPLC) analysis proved that MC production was entirely abolished in Δ*mcyB* strains (Fig. S1B). The cells were grown in BG11 medium at 25 °C under 15 μmol photons m^−2^ s^−1^. Before transfer into the nitrate-free medium, exponential cells grown under N-sufficient conditions were harvested by centrifugation at 6000 *g* for 5 min. The samples were washed three times with nitrate-free medium and then adjusted to the concentration of 1 × 10^7^ cells mL^-1^. Recovery experiments were initiated by adding sodium nitrate solution into the growth medium. The growth of each sample was determined by the Multisizer 3 coulter counter (Beckman Coulter, USA), and the absorption spectra of cells were measured using a UV-VIS spectrophotometer SPECORD^®^ 210 PLUS with the aid of Integration skugel SPECORD (S/N: 25OBO89, Analytik Jena, Germany).

### Determination of MC content

MC extraction was performed as previously described with some modifications (29). About 30 mL of *Microcystis* cells were collected by centrifugation and dried at 40 °C in an oven. The cells were then resuspended in 80% methanol and broken by ultrasonication. The debris and unbroken cells were removed by centrifugation, and the supernatants were collected, evaporated at 40 °C to dryness, and then reconstituted in Milli-Q water (18.2 MΩ cm^−1^). The MC extracts were further purified by a solid phase extraction column (Navigator, China). After stringent washing with 10% methanol to omit nonspecific binding, MCs were eluted with a methanol solution containing 0.1% (v/v) trifluoroacetic acid. MCs were detected and quantified by a LC1200 HPLC system (Agilent, USA) equipped with an XBridge C18 column (4.6 × 250 mm, 5 μm, Waters, USA). The mobile phase used a gradient elution program with a mixture of 0.1% trifluoroacetic acid in water (mobile phase A) and methanol (mobile phase B), respectively. The flow rate was 0.8 mL min^−1^, and the injection volume was 50 μL. The analysis was performed as described by Purdie et al. (30). The detection wavelength was 238 nm. The results were normalized to cell number.

### Chlorophyll fluorescence measurements

Chlorophyll fluorescence measurements were performed with a Joliot-type spectrometer (JTS-10, BioLogic, France) as described previously (31). The maximal PSII quantum yield was measured and calculated as Fv/Fm. The maximal activity of PSI was determined as light-induced P_700_ photo-oxidation in the presence of 10 μM 3-(3,4-dichlorophenyl)-1,1-dimethylurea (DCMU) and 10 μM 2,5-dibromo-3-methyl-6-isopropylbenzoquinone (DBMIB), which block electron transfer from both PSII and cytochrome *b*_6_*f*. The results were normalized to the cell number.

### Flow cytometry analysis

Flow cytometry analysis was performed as previously described (32). Briefly, samples from each time point were collected and fixed with glutaraldehyde (Sigma-Aldrich) at a final concentration of 0.2%. The fixed samples were frozen in liquid nitrogen and stored at −80 °C. The change of auto-fluorescence intensity of *Microcystis* cells was analyzed by a BD FACSVerse flow cytometer (BD Biosciences) with an excitation wavelength of 488 nm and an emission wavelength of 700 ± 54 nm. Data analysis was performed using FlowJo software (v10.8.1).

### Transmission electron microscopy

*Microcystis* cells were harvested by centrifugation at 8000 *g* for 5 min, re-suspended in phosphate buffer containing 2.5% glutaraldehyde, and incubated at 4 °C overnight. After centrifugation, the cell pellets were washed twice with fresh medium and then fixed in a 1% (w/v) potassium permanganate solution. Sample preparation was performed as previously described (33). Ultrathin sections were cut to a thickness of 74 nm using an ultramicrotome (Leica EM UC7, Germany) and stained with 3% uranyl acetate in a lead citrate solution. Images were acquired using a HT-7700 transmission electron microscopy (TEM, Hitachi, Japan) operated at 120 kV.

### Protein preparation and proteomic analysis

Triplicate cultures of *Microcystis* from each time point were collected for proteomic analysis. Proteins were extracted, quantified, and digested according to the method described by Nimer et al. (34). The samples were reconstituted with mobile phase A (2% acetonitrile, 0.1% formic acid), centrifuged at 20,000 *g* for 10 min, and the supernatant was taken for loading. Separation was carried out by a LC-20AB liquid system (Shimadzu, Japan) equipped with a 5 µm, 4.6 × 250 mm Gemini C18 column (Phenomenex, USA) at a flow rate of 300 nL min^−1^. After being separated by a non-linear gradient of mobile phase B (98% acetonitrile, 0.1% formic acid), the end of the nanoliter liquid phase separation was directly connected to the timsTOF Pro spectrometer (Bruker, Germany) for mass spectral analysis. The MS/MS spectra were searched against the protein database of *Microcystis aeruginosa* PCC 7806 in NCBI (https://www.ncbi.nlm.nih.gov/assembly/GCA_002095975.1) using the Maxquant software (v1.6.2.0). The false discovery rate was set to 0.05 for protein identification. Differentially abundant proteins were analyzed by MSstats (v3.22.1), according to a fold change > 2 and adjusted *p*-value < 0.05 as the screening criteria.

### RNA preparation and transcriptomic analysis

Triplicate cultures of *Microcystis* from the indicative growth phase were collected and quickly frozen in liquid nitrogen. Total RNA was extracted using the RiboPure-Bacteria Kit (Applied Biosystems, USA) according to the manufacturer’s instructions. The RNA samples were then quantified using NanoDrop and Agilent 2100 bioanalyzer (Thermo Fisher Scientific, USA), and the samples were sent to BGI Genomics Co. Ltd. for library preparation and RNA sequencing. Bioinformatics analysis was performed using a cloud platform (https://report.bgi.com/ps/mrna/index.html) based on the data generated by the Illumina platform. The expression level of genes was quantified by RSEM software (v1.2.8). Differentially expressed genes were identified by using the DESeq package. The complete transcriptome results are provided in the Supplemental Data files.

## RESULTS

### Toxic *Microcystis* is better adapted to long-term nitrogen starvation

To probe the physiological changes of toxic and non-toxic *Microcystis* under long-term N starvation, *Microcystis* 7806 and its MC biosynthesis deficient mutant (*ΔmcyB*, Fig. S1) were grown in a combined-nitrogen free medium for a period of 30 days (Fig. 1A). Interestingly, unlike the quick bleach of *Synechococcus* sp. PCC 7942 and *Synechocystis* sp. PCC 6803 cultured under low N stresses (35, 36), *Microcystis* 7806 maintains some extent of blue-green color even after prolonged exposure to N starvation (Fig. 1B). This indicates the unique adaptive strategy of this species is slightly different from other unicellular freshwater cyanobacteria. Although *ΔmcyB* (non-toxic) strains showed a resembling growth pattern to the wide type (toxic strain) under nitrogen-sufficient conditions (Fig. S2), their growth under N deficiency was significantly reduced (Fig. 1A). After 30 days of growth in the N-deplete medium, the cell number of the non-toxic strain was approximately 81% that of their toxic homologue.

**FIG 1.**
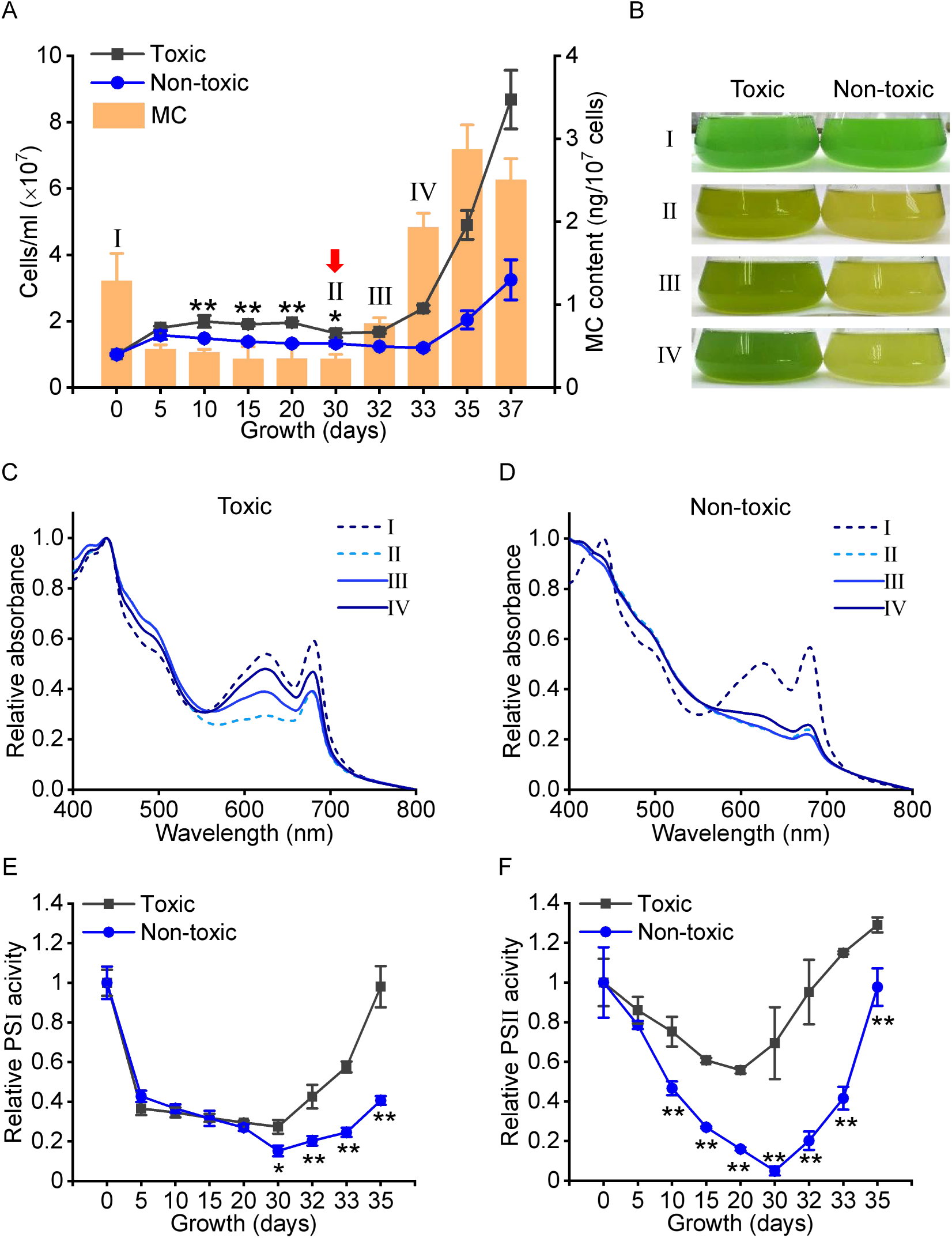
Physiological changes of toxic and non-toxic *Microcystis* during long-term nitrogen starvation and subsequent recovery. (A) The growth curve and intracellular MC content of *Microcystis* strains under different N availabilities. Values are mean ± SD of three independent biological replicates. The results of MC content were normalized to the cell number. Roman numerals mark different phases of the growth experiment, and the arrows mark the days when the recovery experiment was initiated. I, the initiation of growth in the nitrate-limited medium; II, *Microcystis* after 30 days of N starvation; III, *Microcystis* after 2 days of nitrate addition; IV, *Microcystis* after 3 days of nitrate addition. (B) Images of the cultures from the indicated growth phase. (C-D) Absorption spectra of *Microcystis* strains from the indicated growth phase. (E) The changes in PSI activities of *Microcystis* cells over the course of N starvation and recovery. The maximum PSI activity was measured as light-induced P_700_ photo-oxidation in the presence of 10 μM DCMU and 10 μM DBMIB. Values are mean ± SD of three independent biological replicates, and the results were normalized to cell number. The 100% values corresponded to the values at day 0 of triplicates. (F) The changes in PSII activities of *Microcystis* cells throughout N starvation and recovery. PSII photochemical potential was measured by chlorophyll fluorescence kinetics and calculated as Fv/Fm. Values are mean ± SD of three independent biological replicates. The 100% values corresponded to the values at day 0. Asterisks indicate a significant difference between toxic and non-toxic strains using the two-sided Student’s *t*-test. *, *p* < 0.05; **, *p* < 0.01.

The absorption spectra of both *Microcystis* strains showed that the absorbance peaks at 625 nm decreased sharply under N starvation (Fig. 1 C and D), which indicates the degradation of phycobiliproteins. After 30 days of growth in the N-deplete medium, no obvious absorbance peak at 625 nm was observed in non-toxic strains (Fig. 1D). However, in the toxic strains, a minor absorption peak was still observed (Fig. 1C). Further analysis shows that the photochemical potential of both photosystems in *Microcystis* strains decreased significantly under N starvation despite the more rapid decrease in photosystem (PS) II activity of the non-toxic strains (Fig. 1 E and F).

We also quantified the intracellular MC content of toxic strains during the period of long-term N starvation. While it has been suggested that N deficiency may induce the expression of MC biosynthetic genes (23), their intracellular MC content decreased significantly as cells proliferated in the combined nitrogen-free medium. The MC content decreased to 36% of the N-replete cells at the beginning 5 days and then remained constant as cell division was restrained by N starvation (Fig. 1A). Overall, these results consistently showed that MC-producing *Microcystis* maintains a higher photosynthetic activity during long-term N starvation.

### Toxic *Microcystis* recovers more quickly from nitrogen starvation

To assess the recovery potential of toxic and non-toxic strains, *Microcystis* exposed to prolonged periods of N starvation were supplied with combined nitrogen (17.6 μM NaNO_3_). Interestingly, the MC biosynthesis recovered immediately in the toxic strain after the addition of nitrate to the medium. The intracellular MC content increased to the nitrate-growth level within 2 days of nitrate addition (Fig. 1A), while at this moment, the absorbance of phycobiliproteins was not fully recovered (Fig. 1C).

As shown by growth curves, after long-term N starvation, the toxic strains could recover and resume cell division within three days of N supplementation (Fig. 1A). In contrast, the restart of the cell cycle in the non-toxic strains was obviously delayed. After three days of nitrate addition, the cell number of the non-toxic strains resembled that under N-starved conditions. The recovery rate of the PSI (Fig. 1E) and PSII (Fig. 1F) activities was also significantly higher in the toxic strains. After being supplied with the combined nitrogen, the absorbance of the toxic strains increased as a function of time at 680 nm (indicative of chlorophyll *a*) and 625 nm (indicative of phycobiliproteins, Fig. 1C). However, similar phenomenon was not observed in the non-toxic strains at the beginning 2 days of the recovery process (Fig. 1D). Unlike that in the toxic strains, no visual color change was observed in non-toxic culture during the first 2 days of recovery (Fig. 1B).

To better understand the differences in the recovery capacity of these two strains, changes in the internal cell structure of *Microcystis* were studied by TEM (Fig. 2). Under N-replete conditions, the thylakoid membranes of *Microcystis* cells were well organized, and no conspicuous difference could be observed between toxic and non-toxic strains (Fig. 2 A and D). Interestingly, unlike the complete absence of thylakoid membranes in chlorotic *Synechocystis* 6803 (36), *Microcystis* 7806 exposed to 30 days of N starvation remained a small amount of poorly structured thylakoid membranes (Fig. 2 B and E). Moreover, the carbon storage polymer polyhydroxybutyrate (PHB), typically observed under N-limitation (36, 37), was accumulated in both toxic and non-toxic strains. Compared to N-starved cells, toxic *Microcystis* showed a reduced PHB granule accumulation after three days of N supplementation (Fig. 2C and Fig. S3). However, the PHB granules in non-toxic strains were still highly accumulated, and their morphology is more similar to that of N-starved samples (Fig. 2F). All these results consistently suggested that toxic *Microcystis* recovered quicker than non-toxic strains after prolonged N starvation.

**FIG 2.**
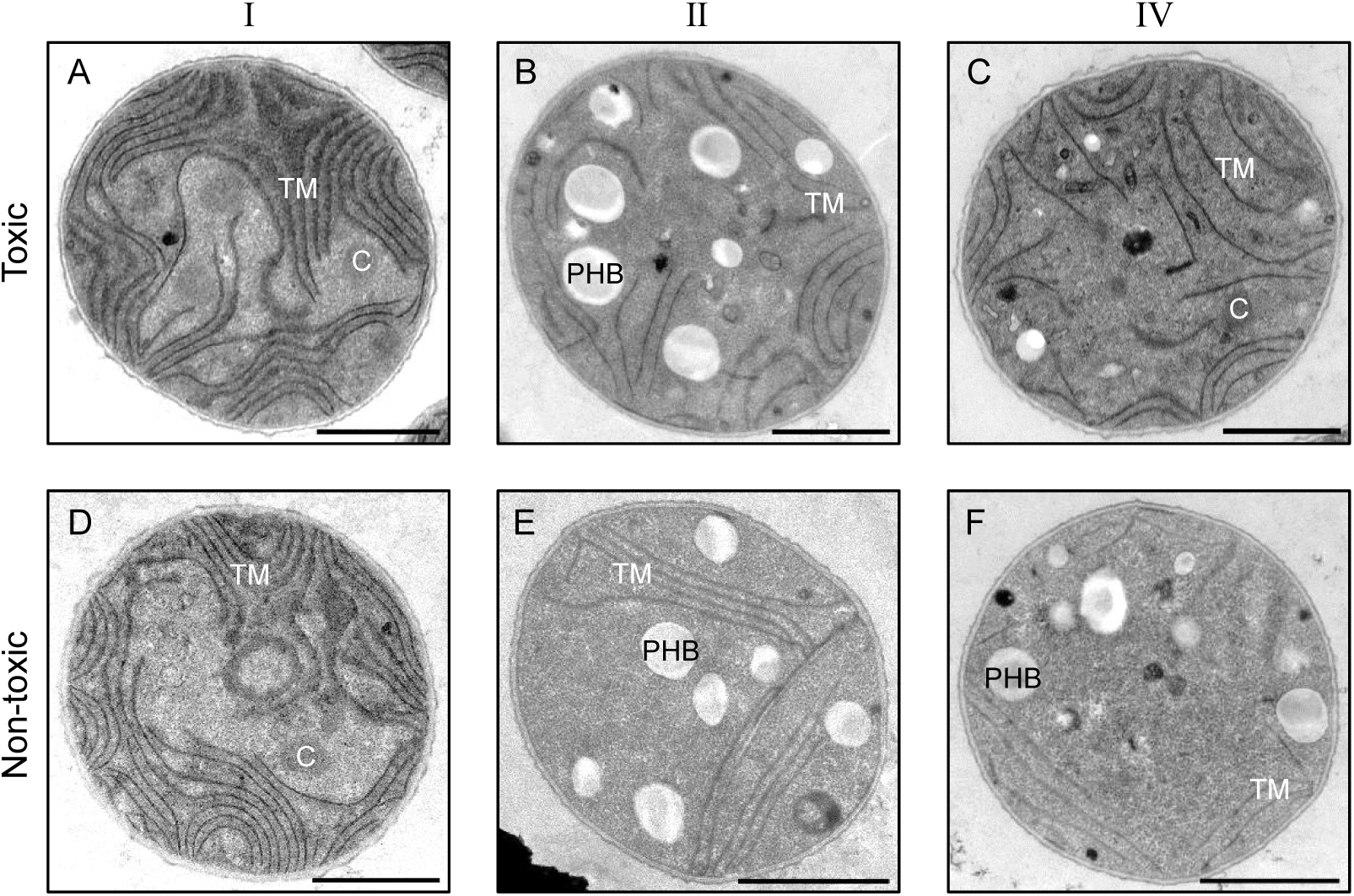
Ultrastructure change of *Microcystis* strains during long-term nitrogen starvation and subsequent recovery. TEM micrographs of the toxic (A-C) and non-toxic (D-F) *Microcystis* cells. (A, D) Cells under N-replete conditions (phase I). (B, E) Cells starved by nitrate for 30 days (phase II). (C, F) Cells on the third day after the addition of nitrate (phase IV). C, carboxysomes; PHB, polyhydroxybutyrate granules; TM, thylakoid membrane. Bar: 1 µm.

### Toxic *Microcystis* is more viable after long-term nitrogen starvation

Previous studies have revealed that most of *Microcystis* 7806 cells were dead under N limitation, and only a small proportion of cells were able to grow and re-green when nitrate was reintroduced (38). To compare the viability of toxic and non-toxic *Microcystis* strains under N fluctuation, we conducted fluorescence-activated cell sorting (FACS) measurements throughout N starvation and recovery processes. FACS analysis shows that the response of cell populations recovering from long-term N starvation was not homogenous (Fig. 3A). Cells exposed to 30 days of N starvation show heterogeneity in the population response to the reintroduction of nitrate into the medium: only part of the population regains chlorophyll auto-fluorescence (high-fl) while the other part of the population remains chlorotic (low-fl). Cell counts of the two subpopulations measured by FACS show that only the high-fl sub-population could grow, whereas the number of low-fl cells remained constant (Fig. 3 B and C). This feature allowed us to estimate the ratio of resuscitable cells. The result shows that the proportion of resuscitable subpopulations from the toxic strains was much higher than that in the non-toxic ones (Fig. 3D), which indicates the high viability of the toxic *Microcystis* under N starvation.

**FIG 3.**
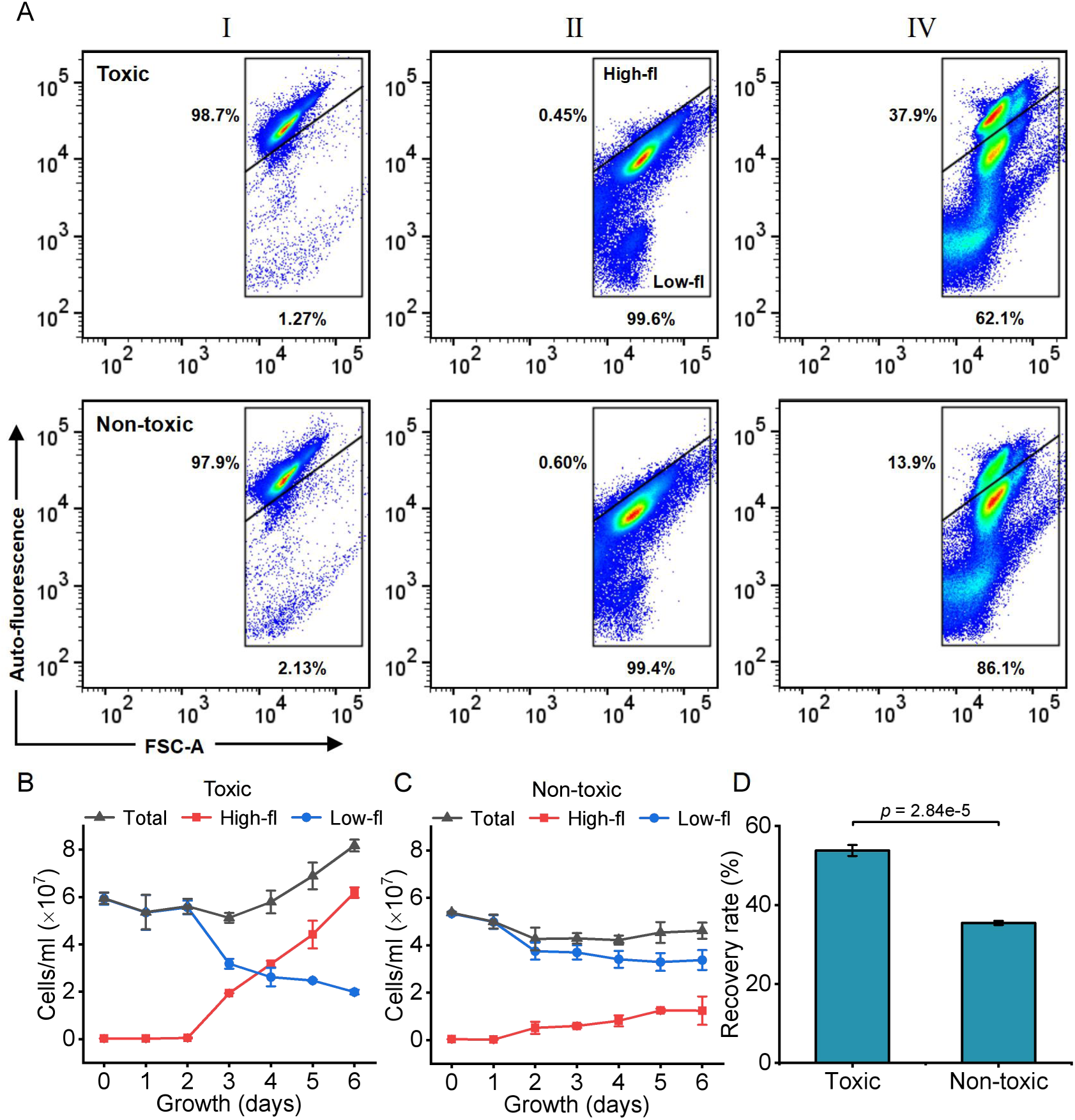
Flow cytometry analysis of *Microcystis* cells during long-term nitrogen starvation and subsequent recovery. (A) 2D scatterplots of samples from growth phases I, II, and IV. The data shown are from a representative sample of three independent biological replicates. FSC, forward scatter; High-fl, high-fluorescence subpopulation; Low-fl, low-fluorescence subpopulation. (B, C) The growth of the cyanobacterial populations of toxic (B) and non-toxic (C) strains during recovery processes. The cell number of *Microcystis* populations was determined before (Total) and after (Highl-fl and Low-fl) gating according to auto-fluorescence intensity. Values are mean ± SD of three independent biological replicates. (D) The proportion of resuscitable cells in toxic and non-toxic strains after prolonged N starvation. The ratio was estimated according to the equation k = 1 − *N_L_*/*N_T_*, where *N_L_* refers to cell numbers of Low-fl subpopulations and *N_T_* refers to the total cell number at the initial of nitrate addition. Values are mean ± SD of three independent biological replicates. Line segments and corresponding *p*-values represent statistical significance calculated by the two-sided Student’s *t*-test.

### Toxic *Microcystis* accumulates more carbon reserves under nitrogen starvation

To gain a deeper insight into the advantages of MC-producing *Microcystis* over non-toxic strains, we analyzed the proteome of both strains after 30 days of N starvation. As shown in Figure 4, most *Microcystis* proteins were more highly expressed in N-starved toxic strains than in their non-toxic homologue. In contrast, only 27 proteins were significantly upregulated in the non-toxic strains (FC > 2, *p* < 0.05, Fig. 4). Amongst them were the permease component of the HCO_3_^−^ uptake system CmpB, subunits of NdhD5/NdhD6 complex, and NhaS3 Na^+^/H^+^antiporter, which belong to the carbon concentrating mechanism (CCM) of cyanobacteria (39).

**FIG 4.**
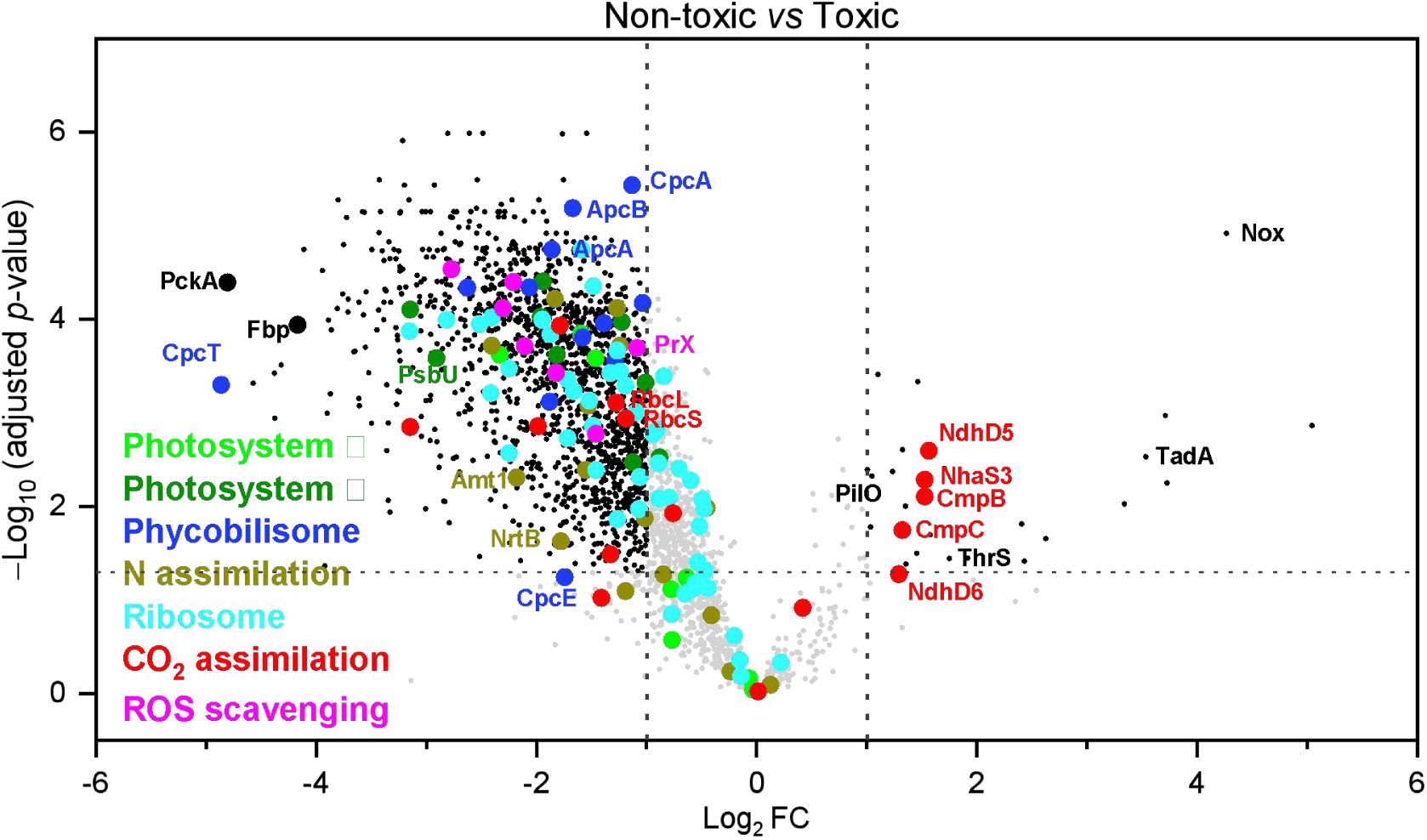
Proteome differences between toxic and non-toxic *Microcystis* after 30 days of nitrogen starvation. Volcano plot showing the global expression pattern of proteins in non-toxic *Microcystis* relative to toxic strains. Broken lines indicate the adjusted *p*-value threshold of 0.05 and Log_2_FC (fold change) thresholds of 1 and −1. Functional groups are color-coded, and proteins without significant differences in expression are shown in light gray. Detailed numeric values are presented in Supplemental Data 1.

Strikingly, proteomic data analysis revealed that two of the most highly expressed proteins in the toxic strains were involved in gluconeogenesis (Fig. 4 and Supplemental Data 1). Among them, phosphoenolpyruvate carboxykinase (PckA) is known to catalyze the conversion of oxaloacetate to phosphoenolpyruvate, whereas fructose-1,6-bisphosphatase (Fbp) is a rate-limiting enzyme in gluconeogenesis, which catalyzes the hydrolyze of fructose-1,6-bisphosphate to fructose-6-phosphate (40, 41). The more highly expressed gluconeogenic genes in toxic *Microcystis* suggested that they might be capable of accumulating more glycogens during N starvation, which could serve as the primary energy source at the initial phase of recovery processes. The quantification of PHB, another important carbon reserve of N-starved cyanobacteria, also supported the increased carbon storage in the toxic strain. After 30 days of N starvation, the toxic *Microcystis* accumulated significantly higher amounts of PHB granules when compared to the non-toxic strains (Fig. S3). All these results consistently show that toxic *Microcystis* accumulates more carbon reserves under nitrogen starvation.

### MC production promotes the resuscitation of nitrogen-starved *Microcystis*

We also probe the metabolic changes of recovering *Microcystis* using a combination of transcriptomic and proteomic analysis. Previous studies in *Synechocystis* have highlighted the importance of respiration at the early phase of recovery from N starvation (36). A similar phenomenon was also observed in the transcriptome of *Microcystis*. Within the first two days of adding the usable nitrogen source, genes involved in Embden-Meyerhof-Parnas (EMP) and oxidative pentose phosphate (OPP) pathways were significantly induced in *Microcystis* (Fig. 5). However, when analyzed the expression of other glycolytic genes, we found that their induction in non-toxic strains is clearly delayed. For example, after three days of recovery from N starvation, only glyceraldehyde 3-phosphate dehydrogenase Gap1 was significantly upregulated at the protein level. In contrast, the expression of other enzymes involved in the glycolysis of non-toxic strains resembled that of N-starved cells (Fig. 5). This upregulated glycogen degradation capacity may contribute to the rapid recovery of toxic strains compared to their non-toxic homologue.

**FIG 5.**
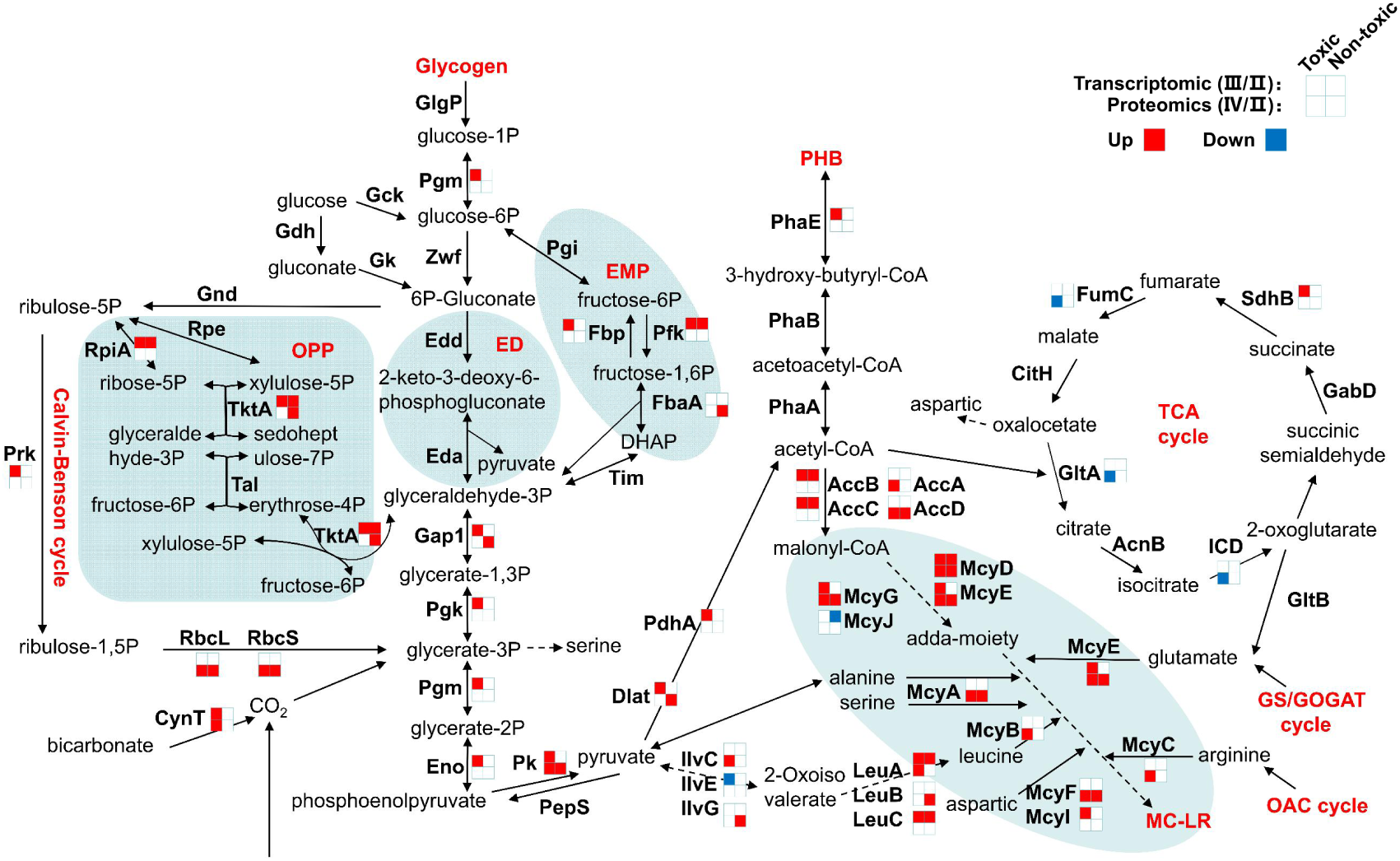
The expression changes of enzymes involved in central carbon metabolism during recovery from nitrogen starvation. The relative change of corresponding enzymes was analyzed using a combination of transcriptomic (phase III) and proteomic (phase IV) analysis, and the result was normalized to N-starved growth (phase II) of each strain. Genes that were differently expressed were marked with red (up-regulated) or blue (down-regulated) code, respectively, whereas those without significant differences in expression were shown in white. The EMP (Embden-Meyerhof-Parnas), ED (Entner-Doudoroff), and OPP (Oxidative Pentose Phosphate) pathways were highlighted by color. The detailed changes of MC synthetic genes are shown in Table S1, and the genome-wide numeric values are presented in Supplemental Data 2.

After three days of recovery from N starvation, the absorbance of phycobiliproteins in toxic *Microcystis* was greatly increased (Fig. 1C), indicating a substantial intracellular N accumulation. Consequently, the depletion of the tricarboxylic acid (TCA) cycle by amino acid synthesis should be alleviated (42). However, our omics data shows that, while glycolytic genes were significantly upregulated in the toxic strains, most genes involved in the TCA cycle were not induced within the first three days of nitrate addition (Fig. 5), indicating a potential risk of acetyl-CoA accumulation. Interestingly, the expression of acetyl-CoA carboxylase genes was found to be upregulated during the recovery processes. Acetyl-CoA carboxylase was known to convert acetyl-CoA to malonyl-CoA, which could serve as the precursor in MC synthesis. Further analysis showed that both transcription and protein levels of MC biosynthetic genes were significantly upregulated in *Microcystis* cells (Fig. 5 and Table S1), suggesting that the MC synthetic pathway resumed at the early resuscitation phase and may serve as an alternative strategy to metabolize the accumulated acetyl-CoA.

After three days of nitrate addition, the expression of genes encoding the subunits of ribulose bisphosphate carboxylase, RbcL and RbcS, was significantly upregulated in *Microcystis* (Fig. 5), which indicated a recovery of photosynthetic carbon sequestration potential. Indeed, many genes involved in tetrapyrrole metabolism (Fig. S4) and photosynthesis (Fig. S5) were upregulated at this time point, which is consistent with our physiological measurements (Fig. 1). Additionally, compared to that of the toxic strains, a delay in the induction of these genes in non-toxic strains could be clearly observed.

## DISCUSSION

Although the ecological importance of harmful cyanobacterial blooms has been long recognized, the genetic traits that provide bloom-forming cyanobacteria a competitive advantage over other species were poorly understood. Several environmental factors were correlated to the development of cyanobacterial blooms, including temperature (5, 43), CO_2_ concentration (44, 45), and N and P availabilities (46, 47). Here, we provided multiple lines of evidence consistently suggesting that MC production is critical for *Microcystis* to survive and recover from long-term N starvation. The presence of MC synthetic genes could enable toxic *Microcystis* to quickly resuscitate and resume growth when conditions are favorable again, which has significant consequences for the composition and succession of *Microcystis* populations.

Unlike the quickly entering the dormant state of many other unicellular non-diazotrophic cyanobacteria (36, 48), *Microcystis* strains maintained vegetative growth even after prolonged N starvation. In this context, how they protect their metabolic activities could be extremely important. While MC synthesis is a high energy and nitrogen cost process, it provides *Microcystis* an advantage under prolonged N starvation. Our research shows that the MC-producing *Microcystis* exhibited a higher resuscitation rate after long-term N starvation. In contrast, the viability of the non-toxic strains was much lower (Fig. 3D). This could be ascribed to the role of MC in protecting against oxidative stress, which suggested that MC could covalently bind to the cysteine residues of Rubisco to avoid conformational changes and enzyme inactivation (49). Our proteomic analysis shows that toxic *Microcystis* experiencing 30 days of N starvation accumulated more RbcL and RbcS proteins than those in non-toxic strains (Fig. 4). Further analysis revealed that proteins more highly expressed in non-toxic strains were enriched in components of CCM. Thus, it is reasonable to speculate that MC-producing *Microcystis* maintains a higher Rubisco activity under N limitation, which helps to accumulate more carbon reserves to support the survival of cells under long-term N starvation and fuel subsequent recovery.

Such a protective role of MC to dark reactions could help to prevent the photo-inhibition of N-starved *Microcystis*. Unlike the complete absence of photosynthetic membranes in the chlorotic *Synechocystis* (36), *Microcystis* exposed to prolonged N starvation still retains a certain amount of thylakoid membrane (Fig. 2). Without the protection of MCs to the Rubisco, the sustained excitation of the photosynthetic apparatus could result in the imbalance between light and dark reactions, which may lead to the photo-inhibition of the non-toxic strains. This hypothesis is further supported by our physiological analysis, as the non-toxic strains exhibit a more seriously declined PSII activity under N starvation when compared to that of the toxic *Microcystis* (Fig. 1F). From this aspect, the upregulation of inorganic carbon transport in non-toxic strains was physiologically necessary, which enable them to quench excess light energy while their carbon sequestration capacities are limited (50–52).

As one of the biggest intracellular N pools, phycobiliproteins were known to be degraded under N starvation and resumed synthesis when provided with usable nitrogen resources (48). Interestingly, our result shows that when nitrate was reintroduced, MC biosynthesis resumed before phycobilisomes were fully recovered, indicating the importance of MC production at the early resuscitation phase. The transcriptomic analysis revealed that, although genes involved in the glycolytic pathway were significantly upregulated in the toxic strains within the first 2 days of nitrate addition, the transcript of TCA cycle genes was not induced coordinately. This would result in a massive accumulation of acetyl-CoA and NADH, which may be inhibiting glycolytic processes (53). Under this background, the initiation of MC production when combined N was available could be extremely important for *Microcystis*, which enabled them to convert metabolic overflow into this stable cyclic heptapeptide on the one hand and inhibit the growth of surrounding eukaryotic and prokaryotic phytoplankton on the other (9, 54).

It should be noted that the ability of *Microcystis* to survive under prolonged N starvation and then recover when conditions are favorable is critical to their survival and success in natural environments, as nitrogen concentration in surface water frequently varies seasonally. Long-term observation revealed that nitrogen concentration in surface water commonly increases in spring and decreases from summer to autumn (27, 28, 46). Interestingly, toxic *Microcystis* was generally dominant at the early stage of bloom formation (55, 56), probably due to their ability to produce MC. Based on our findings, a hypothesized model can be proposed (Fig. 6). *Microcystis* were known to overwinter on the sediment surface and serve as an inoculum for the following year’s epilimnetic growth (57). When temperature and N availability are increased, *Microcystis* cells recover their activities, and the MC-production may provide toxic *Microcystis* a competitive advantage over non-toxic strains (19), which enables them to quickly resuscitate in early spring and become the dominant species during bloom development. While the quick consumption of N resources during dense blooms is not conducive to MC synthesis, the decreased temperature in early winter may stimulate MC accumulation under low N conditions (47, 58), which may help toxic *Microcystis* to survive during prolonged N-starved winter.

**FIG 6.**
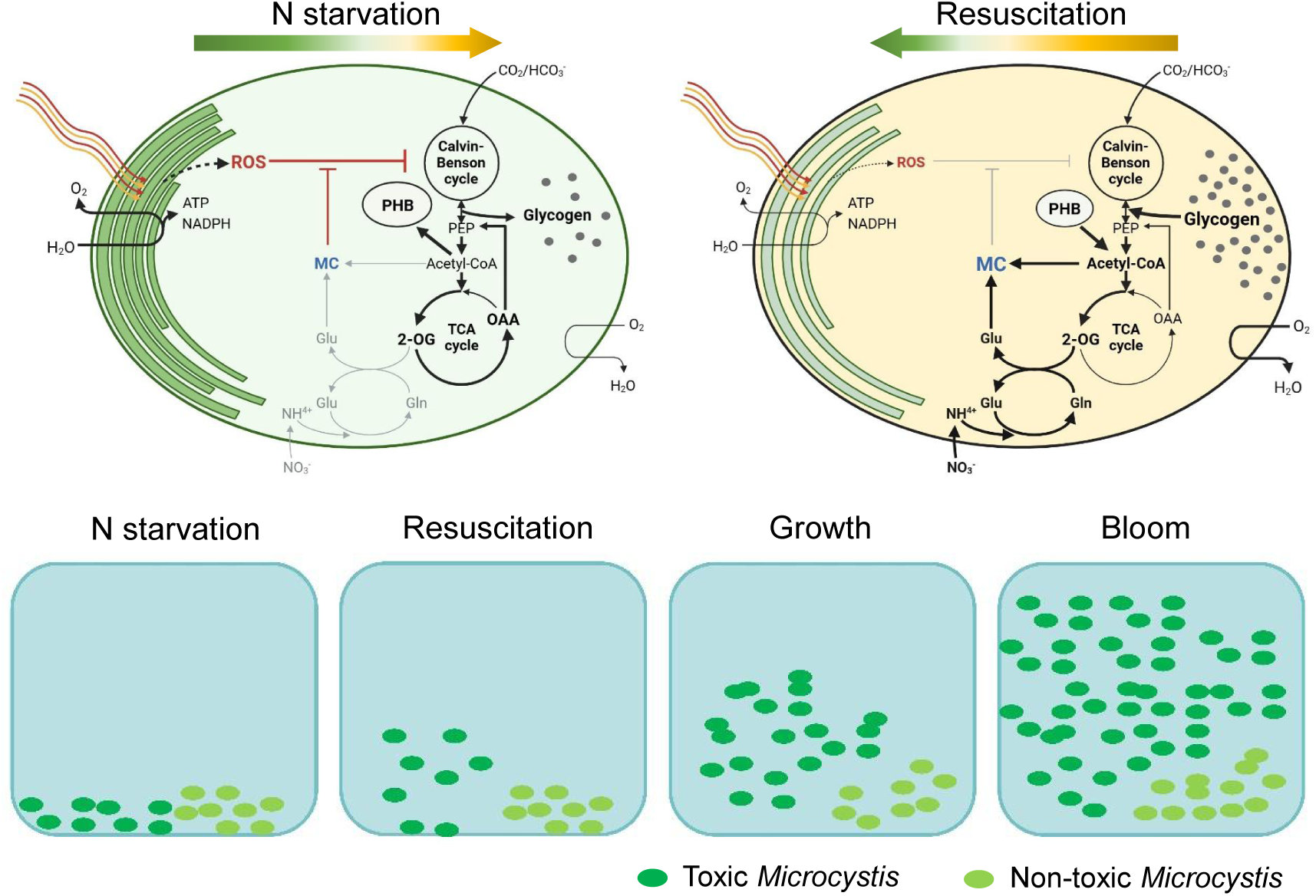
Schematic illustration of the MC-mediated N fluctuation acclimation pathways in *Microcystis*. The metabolic pathways and reactions up-regulated under indicated conditions are shown in bold arrows. The figure was created with BioRender.com.

While the *Microcystis* genus is characterized by its high morphological diversity, phylogenetic analysis revealed that all *Microcystis* species whose genomes or 16S rRNAs have been sequenced warrant placement into the same species complex (59). This observation demonstrates the close genetic traits between *Microcystis* strains and suggests the strategies we observed might be highly conserved. Our results show that MC production is critical to maintaining the carbon metabolism of toxic *Microcystis* during long-term N starvation and subsequent recovery. This could be extremely important for understanding the population succession of cyanobacterial blooms, as increased N loading and high N:P ratio have made freshwater blooms more toxic (47).

## ACKNOWLEDGEMENTS

We thank Prof. Cheng-Cai Zhang (Institute of Hydrobiology, Chinese Academy of Sciences) for his valuable advice. We also thank the staff from the Analysis and Testing Center of Institute of Hydrobiology, Chinese Academy of Sciences for the help with TEM microscopy operation.

## FUNDING

This work was supported by the National Key R&D Program of China (2018YFA0903100) and National Natural Science Foundation of China (92051105, 32270397).

## AUTHOR CONTRIBUTIONS

Xiao-Ya Lian, Zhong-Chun Zhang, and Bao-Sheng Qiu designed the research; Xiao-Ya Lian, Guo-Wei Qiu, Wen-Can Zheng, and Jin-Long Shang carried out the experiments; Xiao-Ya Lian, Guo-Wei Qiu, Hai-Feng Xu, Guo-Zheng Dai, Nan-Qin Gan, Zhong-Chun Zhang, and Bao-Sheng Qiu analyzed the data; Xiao-Ya Lian, Guo-Wei Qiu, Zhong-Chun Zhang, and Bao-Sheng Qiu wrote the manuscript.

## DATA AVAILABILITY STATEMENT

The transcriptomic data generated in this study have been deposited in NCBI SRA database under the accession code PRJNA1115087: https://www.ncbi.nlm.nih.gov/bioproject/PRJNA1115087. The proteomics data have been deposited to the ProteomeXchange Consortium under accession code PXD053031: https://proteomecentral.proteomexchange.org/cgi/GetDataset?ID=PXD053031.

## CONFLICT OF INTEREST

The authors declare no conflicts of interest.

## Supplemental Material

**Supplemental material.** Figs. S1 to S5; Table S1.

**Supplemental Data 1.** The complete list of the differentially expressed proteins in toxic and non-toxic *Microcystis* after 30 days of nitrogen starvation.

**Supplemental Data 2.** The transcriptome and proteome differences of *Microcystis* strains over the time course of recovery.

## Notes

### Competing Interest Statement

The authors have declared no competing interest.

