## Supplemental material for "Microcystins are critical for the toxic *Microcystis* to survive long-term nitrogen starvation"

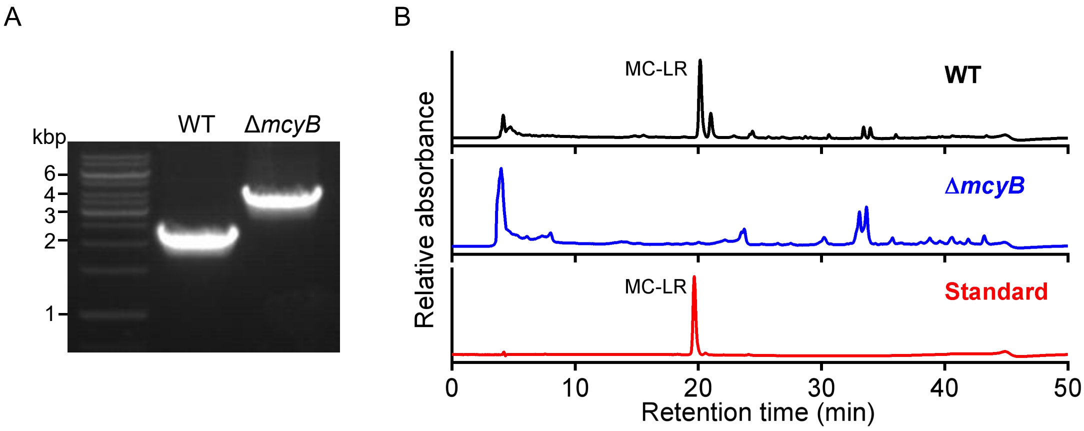


**FIG S1** Characterization of Δ*mcyB* strain. (A) PCR results demonstrated that the chloramphenicol-resistant fragment was successfully inserted into the open reading frame of the *mcyB* gene. (B) HPLC analysis confirmed the MC production was entirely abolished in Δ*mcy*B strains. Similar results were obtained from three independent experiments.

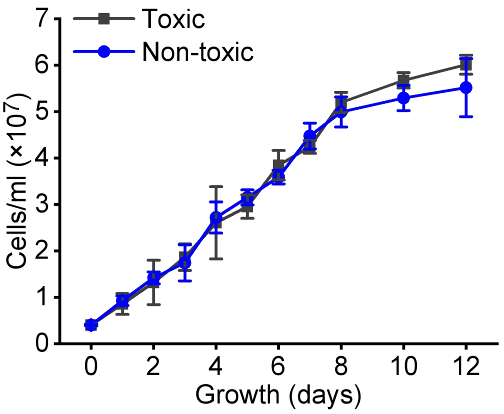


**FIG S2** The growth of toxic and non-toxic *Microcystis* under nitrate-replete conditions. Values are mean ± SD of three independent biological replicates.

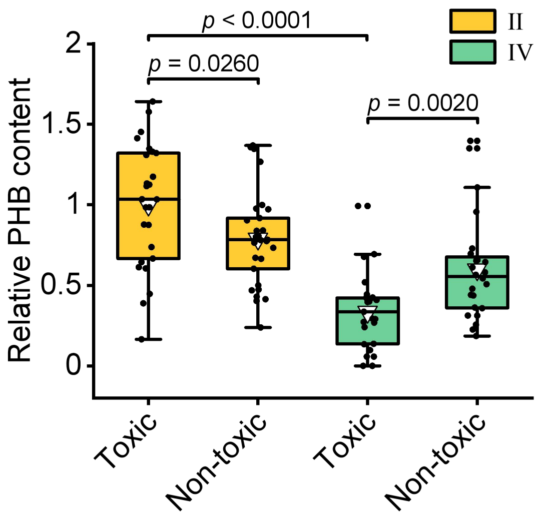


**FIG S3** Change in PHB content during recovery from N starvation. The relative PHB content of *Microcystis* cells was estimated from the area of polyhydroxybutyrate granules under transmission electron microscopy. Data shown were measured with ImageJ software from at least 24 cells, and the results were normalized to that of the N-starved toxic strains. The embedded line and triangle in the boxplot mark the median and mean values, respectively. The upper and lower edges of the box indicate the first and third quartiles, and the whiskers extend to 1.5× the interquartile range beyond the first and third quartiles. Line segments and corresponding *p*-values represent statistical significance calculated by the Mann-Whitney test.

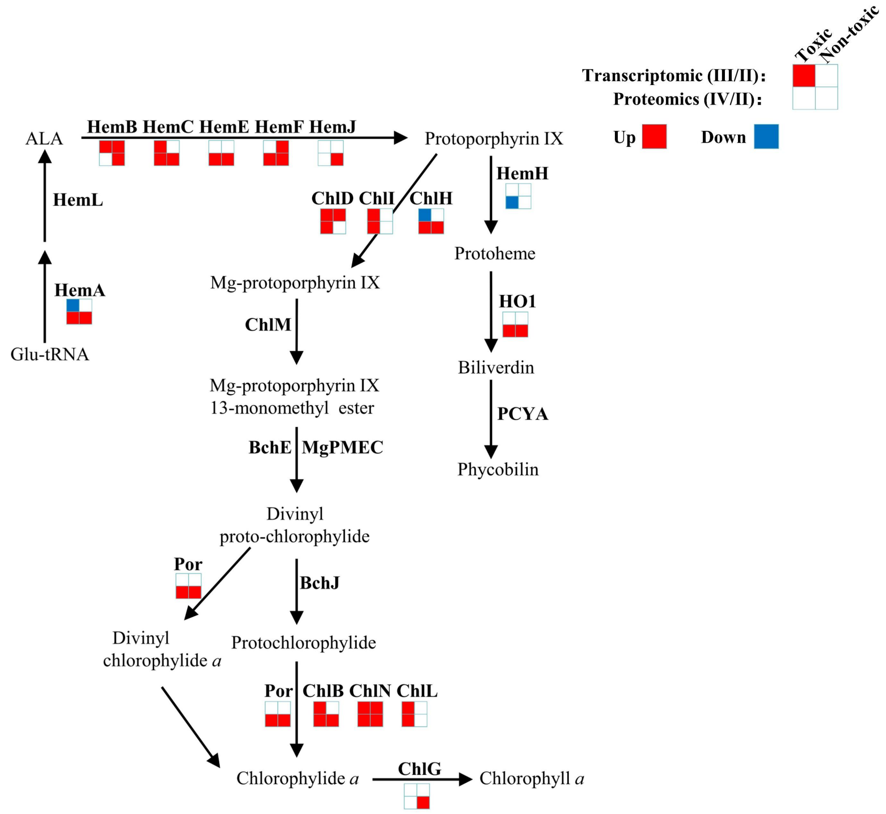


**FIG S4** The change of tetrapyrrole metabolic genes during recovery from N starvation. The relative expression change of corresponding genes was analyzed using a combination of transcriptomic (phase III) and proteomic (phase IV) analysis, and the result was normalized to N-starved growth (phase II) of each strain. Genes that were differently expressed were marked with red (up-regulated) or blue (down-regulated) code, respectively, whereas those without significant differences in expression were shown in white. The genome-wide numeric values are presented in Supplemental Data 2.

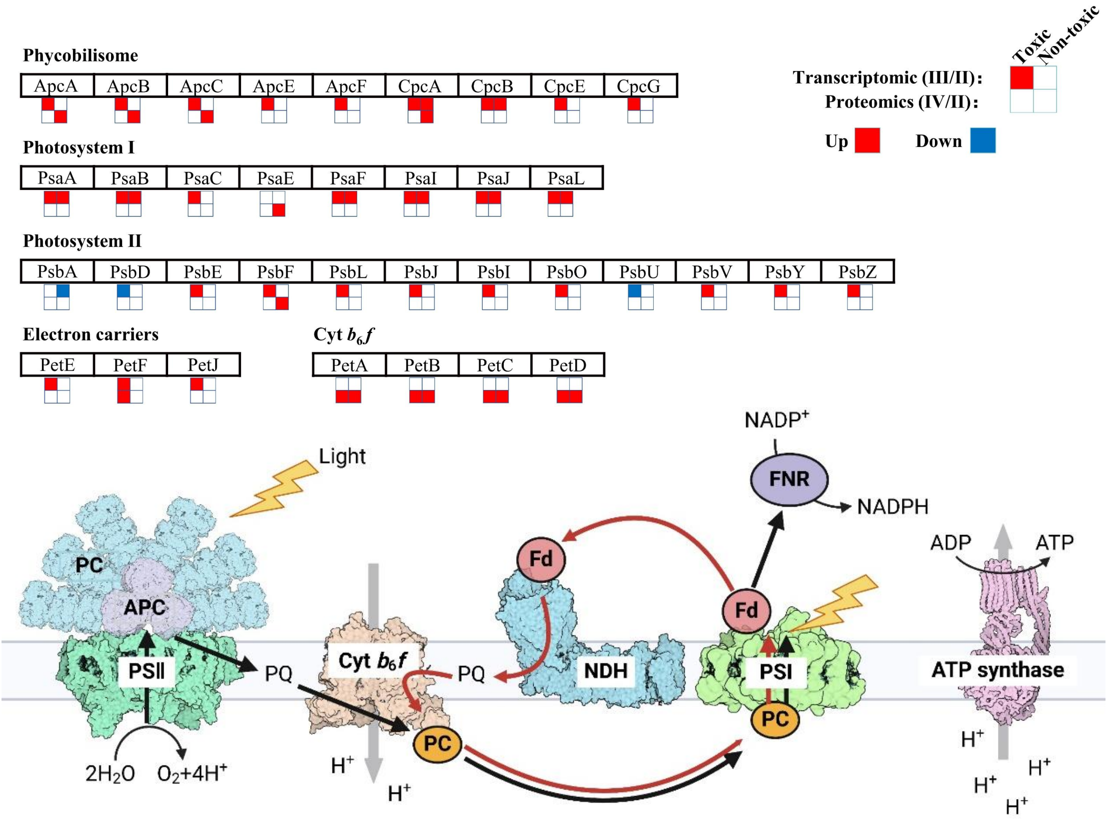


**FIG S5** The change of photosynthetic genes during recovery from N starvation. The relative expression change of corresponding genes was analyzed using a combination of transcriptomic (phase III) and proteomic (phase IV) analysis, and the result was normalized to N-starved growth (phase II) of each strain. Genes that were differently expressed were marked with red (up-regulated) or blue (down-regulated) code, respectively, whereas those without significant differences in expression were shown in white. The genome-wide numeric values are presented in Supplemental Data 2. The figure was created with BioRender.com.

**TABLE S1** Change of MC synthetic genes during recovery from N starvation.

| Gene | Transcriptomic III/II | | Proteomics IV/II | |
|---|---|---|---|---|
| | $Log_2FC$ | *p*-value | $Log_2FC$ | *p*-value |
| *mcyA* | − | − | 2.9912 | 6.09E-07 |
| *mcyB* | − | − | 1.1248 | 4.31E-05 |
| *mcyC* | − | − | 1.1931 | 3.21E-03 |
| *mcyD* | 1.9179 | 4.67E-06 | 2.1637 | 7.76E-04 |
| *mcyE* | 1.1867 | 4.96E-06 | 1.5366 | 1.62E-05 |
| *mcyF* | 1.2779 | 1.28E-04 | − | − |
| *mcyG* | 1.2470 | 1.71E-04 | 2.1750 | 1.13E-04 |
| *mcyI* | − | − | 1.7479 | 1.45E-04 |
| *accA* | − | − | 1.9284 | 8.83E-06 |
| *accB* | 1.2777 | 3.66E-07 | − | − |
| *accC* | 2.7774 | 6.19E-11 | − | − |
| *accD* | − | − | 1.2841 | 2.45E-03 |
